# Towards unsupervised classification of macromolecular complexes in cryo electron tomography: challenges and opportunities

**DOI:** 10.1101/2022.03.10.483789

**Authors:** E. Moebel, C. Kervrann

## Abstract

**Background and Objectives:** Cryo electron tomography visualizes native cells at nanometer resolution, but analysis is challenged by noise and artifacts. Recently, supervised deep learning methods have been applied to decipher the 3D spatial distribution of macromolecules. However, in order to discover unknown objects, unsupervised classification techniques are necessary. In this paper, we provide an overview of unsupervised deep learning techniques, discuss the challenges to analyze cryo-ET data, and provide a proof-of-concept on real data.

**Methods:** We propose an unsupervised sub-tomogram classification method based on transfer learning. We use a deep neural network to learn a clustering friendly representation able to capture 3D shapes in the presence of noise and artifacts. This representation is learned here from a synthetic data set.

**Results:** We show that when applying k-means clustering given a learning-based representation, it becomes possible to satisfyingly classify real sub-tomograms according to structural similarity. It is worth noting that no manual annotation is used for performing classification.

**Conclusions:** We describe the advantages and limitations of our proof-of-concept and raise several perspectives for improving classification performance.

## 1 Introduction

Cryo electron tomography (cryo-ET) enables to visualize sub-cellular environment at nanometer scale, allowing to identify macromolecular complexes in their native state. The field of view and the resolution are both large enough to enable a joint study of the cellular context and structures. Therefore cryo-ET acts as a link between low resolution (e.g., light microscopy) and high resolution (e.g., X-ray crystallography) imaging techniques, filling the gap of knowledge between several scales. However, noise and artifacts in cryo-ET are such that heavy computational processing is needed to access the image content.

In recent years, a number of supervised deep learning based techniques have been developed for identifying macromolecular species in cellular cryo-ET [1, 2]. Although effective, the application of supervised methods to cryo-ET still remains limited for several reasons. First, a significant amount of annotated data is required to train deep neural networks. Unfortunately, manual and semi-automatic annotation is a time consuming and tedious task, and prone to bias. Second, the generalization is not guaranteed when an input tomogram has been acquired with different parameters and imaging conditions compared to training tomograms. While preliminary results [2] exist, transferring learning across experimental contexts is an open question. Finally, biologists are bound to identifying already known macromolecule species. Identifying new and unknown species (or conformational states of a species) requires unsupervised classification techniques as already investigated in [3] [4] [5].

In the remainder of the paper, we present in Section 2 an overview of unsupervised deep learning methods and their potential application to cryo-ET. In Section 3, we describe our proof-of-concept method, which was especially designed to cope with the limitations imposed by cryo-ET data. In Section 4, we present preliminary results showcasing the potential of our approach. Finally, in Section 5 we discuss these results, and propose several ideas to improve the performance obtained so far.

## 2 Unsupervised deep learning and its promises in cryo-ET

### Unsupervised deep learning

This research field is very active in artificial intelligence (AI) and encompasses several topics, such as representation learning, self-supervised learning, clustering, as well as transfer learning. We detail below these different concepts and explain how they relate to each other.

Deep neural networks learn from data through the optimization of an objective cost function. The optimization algorithm is exclusively of the family of stochastic gradient descent. Therefore, the necessary condition to employ deep neural networks, regardless of the setting (supervised or unsupervised), is the ability to generate a significant gradient signal. This is achieved by designing a loss function which evaluates the distance of the network output to a so-called training target. In the supervised setting this target is a set of annotations (i.e., labels). In the unsupervised setting, the target is either a data model or is obtained via self-supervised learning. In the self-supervised setting, the target is generated automatically from the data using ad-hoc heuristics. For instance, auto-encoders [6] are typical self-supervised learning methods, for which the targets are identical to the input data of the network. Here, the goal is not to classify data, but to determine low-dimensional and reversible representations. In the literature, a variety of self-supervised tasks have been proposed for image representation and analysis, such as in-painting [7], colorization [8] or predicting transformations [9]. The fact that humans are not involved in the target generation is why deep neural networks are considered to achieve unsupervised learning.

### Representation learning

This process [10] allows us to learn a (non-linear) mapping between the raw data space and an alternate feature space, in which the task at hand is easier to solve. The feature space is a low dimensional representation of raw data and encodes more abstract concepts, which tends to boost the performance on downstream tasks. If the downstream task is clustering, as it is the case for unsupervised classification, then it is desirable that data points are separable with respect to the characteristics of interest. The representation learning task can be either supervised (e.g., transfer learning) or self-supervised (e.g., contrastive learning, deep clustering).

### Transfer learning

In this approach [11], the representation learned for condition *A* is used for condition *B*. The underlying assumption is that the same representation may be useful in both conditions. This concept is well understood in supervised image classification, and arises from the observation that the first layers of CNNs (near the input) tend to resemble Gabor filters or color blobs, regardless of the task at hand. It is well established [11] that the first layers capture low-level features such as edges and shapes, and are relevant for tasks as diverse as dog breeds and mushroom species classification.

A useful application of this feature transferability is when the available training set is small and the risk of over- or underfitting exists. In this case, one can benefit from networks trained on a huge data sets (e.g., ImageNet [12]), by *freezing* the weight-values of the first layers and therefore train only the last task-specific layers. In this way only few weights have to be trained and computation is less demanding. Another strategy is to apply *fine-tuning*; instead of freezing transferable weights, they are used to initialize a network. The consequence is that the network parameters are closer to the target configuration (compared to random initialization), therefore significantly accelerating the convergence of the training procedure.

Transfer learning is also used to acquire general purpose feature extractors (i.e., deep features). It is now common to re-use as a representation the output of intermediate layers of networks that have been trained on ImageNet (e.g. VGG or ResNet). These representations are exploited to solve various problems, including unsupervised classification. As described in [13], coupling clustering algorithms such as k-means with these representations can lead to remarkable classification accuracies. That being said, while representations derived from ImageNet are very powerful for natural images (i.e., photographs), it is not always the case when they are applied to other imaging modalities like microscopy [14], as illustrated and discussed later on. Depending on the problem, these representations may not discriminate the desired features.

### Contrastive learning

These methods [15] have received interest due to success in self-supervised representation learning. The strategy relies on comparison of sample pairs, instead of individual samples. The idea being that, in the learned representation, similar samples should be mapped together, while dissimilar samples should be pushed away. The key issue is to define pairs of similar and dissimilar samples. This is resolved by applying strong image augmentation (i.e., transformations like rotation, cropping, resizing, blur etc). Similar pairs are then obtained as being distorted copies of the same sample, and dissimilar pairs as being different samples.

In the case of cryo-ET data, care must be taken as to which transformations to use. In fact, in a macromolecule classification task, the main clue is its structure (including size and chirality). Therefore transformations such as resizing, mirror operations or elastic deformations (i.e., diffeomorphism) are not appropriate. In addition, the missing wedge is responsible for anisotropic resolution in the 3D image, inducing delocalization of densities along the *z* direction, as well as loss of information in the *xy* plane along the direction perpendicular to the tilt axis. We do not believe that a network should be invariant to the orientation of this anisotropy, and hence using random 3D rotations for data augmentation should be avoided. This leaves us with few choices left, like random translations and linear transformations on voxel values (e.g., contrast changes), which limits the potential of contrastive learning methods in cryo-ET.

### Joint representation learning and clustering

In order to guaranty that a learned representation is clustering friendly, a family of methods named deep clustering [16, 17, 18] adapts the underlying training objective. Here, the training loss is a combination of self-supervised and clustering-specific objectives. The self-supervised objective is used to impose constraints on the learned representation, and popular choices include the reconstruction loss or the self-augmentation loss [16] (derived from contrastive learning). The clustering-specific objective ensures that the learned representation is clustering-friendly, and includes losses such as the k-means loss, cluster assignment hardening, and group sparsity loss [16]. The clustering itself is generally performed by k-means (centroid-based) or agglomerative clustering (hierarchical).

Nowadays, several unsupervised deep learning methods achieve promising accuracies on benchmark data sets such as MNIST (*>* 95%) and ImageNet (*>* 80%), reducing the performance gap with supervised methods. This being said, supervised methods will always perform better, as confirmed by experiments on these benchmarks.

### Application to cryo-ET

While these performances are encouraging and represent good opportunities in cryo-ET in the future, the aforementioned methods provided results in well-defined conditions. The conventional benchmarks constitute an ideal classification setting: the data is not corrupted by noise, the classes are equally balanced (same number of examples per class), and the data set size is huge (ImageNet consists of more than 1 million images). These properties are not satisfied with cryo-ET data. Here, the SNR level is exceptionally low, and tomograms are corrupted by imaging artifacts (e.g., missing wedge). Macromolecule populations vary significantly, and the target species may be rare. Finally, the orders of magnitude of dataset size differ: for cryo-ET, a data set composed of 50 tomograms containing 10000 instances of a macromolecule class (e.g., ribosomes) is considered as large. But more frequently, datasets are constituted of approximately 10 tomograms.

As shown in [14], the performance of deep clustering methods depends on the amount, quality, and type of the input data. The authors show that among tested data types, bioimage data was the one for which these methods perform worst, and conclude that these methods are ”*not suitable for small-sized data sets*”. Furthermore, it is not clear how these techniques behave when a class is rare: will clusters be detected if the number of members is too low?

When considering an application such as cryo-ET, an additional limitation of deep learning-based representation learning is interpretability. There is no direct way of identifying which attributes the classification is based on, which is a major disadvantage for gaining insights on detected clusters.

In this paper, we present a proof-of-concept of what appears to us the most sensible choice for achieving unsupervised classification in cryo-ET. We propose to use a synthetic cryo-ET data set (i.e., a data model) to learn a clustering-friendly representation. This learned representation is then used for clustering experimental data. This approach has several advantages. First, the learned representation is interpretable, because we define explicitly the training target (i.e., the labels) and therefore we control what the network encodes. Secondly, this strategy is scalable, as we can generate as much synthetic training data as needed. This makes the approach appealing to analyze both small and large data sets (i.e., real tomograms). Third, this approach is adapted for discovering rare macromolecule species, as the representation learning does not depend on manual annotation. Therefore it is not affected by the issue of under-represented classes, because training data generation is essentially free. We illustrate this concept in the setting of subtomogram classification (which can be extended to segmentation).

## 3 A proof-of-concept method for unsupervised low-dimensional representation learning

Our objective is to find a non-linear mapping *f*_*θ*_ : *X* → *Z* where *X* ∈ ℝ^*D*^ is the data space and *Z* ∈ ℝ^*N*^ is the learned representation, where *N < D*. This representation should (i) satisfyingly characterize 3D shapes in the presence of noise and artifacts, as observed in cryo tomograms. It should also be (ii) rotation invariant, in order to avoid subtomogram alignment as in [4] and [5]. Once such a representation has been derived, one can apply classical clustering algorithms (e.g., k-means) on these feature vectors to group unknown objects according to their structural similarity (see Fig. 1 B). Consequently, applying subtomogram averaging to objects belonging to the same cluster should reveal the high-resolution structure of unknown objects. As we will see, in our case no clustering loss was necessary to yield good clustering results.

**Figure 1:**
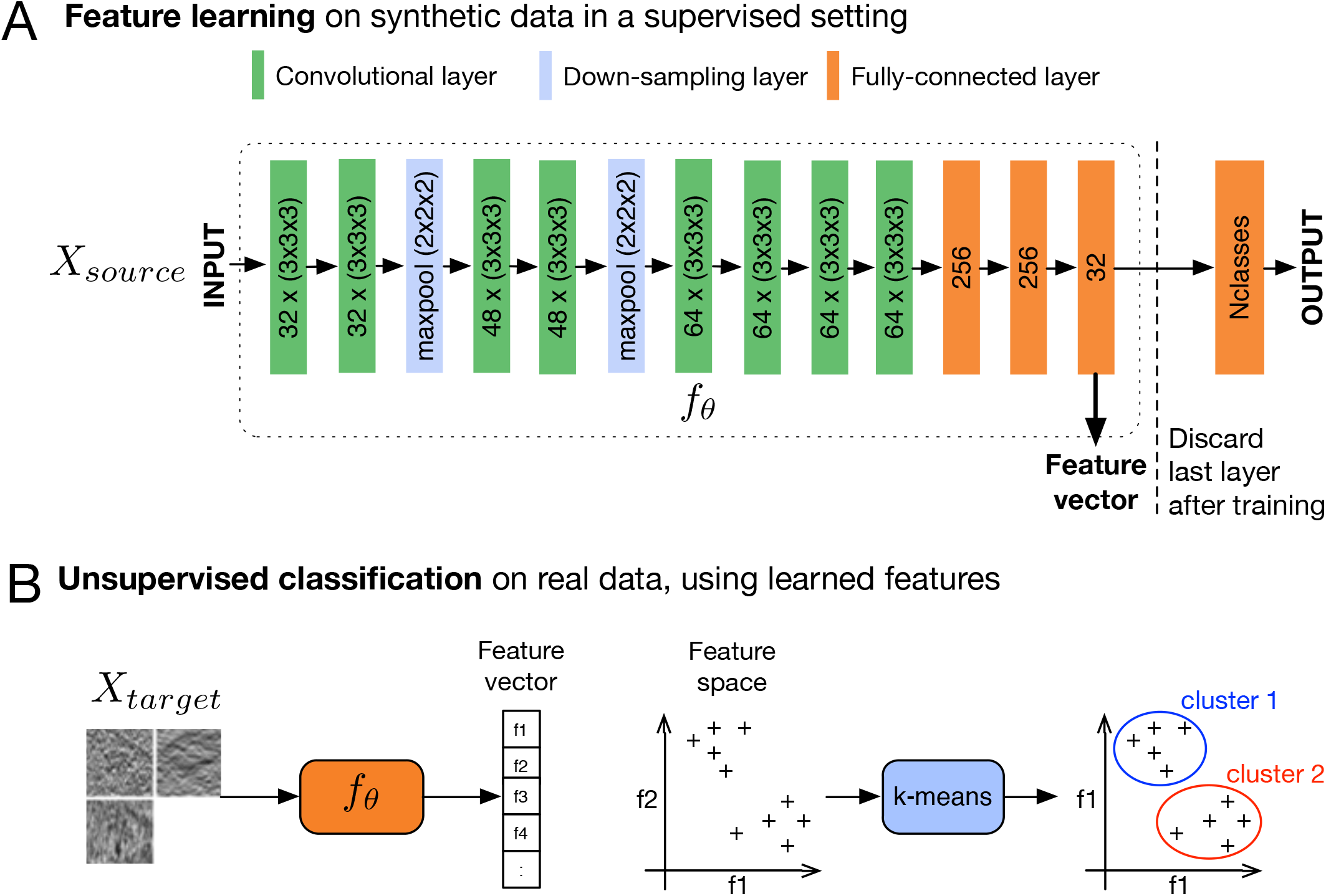
Unsupervised subtomogram classification. (A) Architecture of the CNN used to learn a feature space that characterizes 3D shapes. The network is trained on a synthetic data-set composed of 13 different macromolecule classes, obtained from the PDB databank. Once trained, we discard the last layer. We now have a network that takes as an input volumes and outputs feature vectors of size 32. (B) We use this network to compute the feature vectors of experimental sub-tomograms. We finally achieve unsupervised classification of these sub-tomograms by applying k-means clustering on their feature vectors. The obtained clusters group the sub-tomograms by structural similarity.

We choose *f*_*θ*_ to be the encoding part of the DeepFinder network from our previous work [2]. The latter is a encoder-decoder architecture designed for segmentation (i.e., Unet like). The parameters *θ* are learned in the frame of a subtomogram classification task. We assume that a mapping *f*_*θ*_ learned from such a task should meet our requirements (i) and (ii) defined above. We finally add fully connected layers as shown in Fig. 1 A to fit the encoder network to the classification task. Once the parameters *θ* have been learned, we obtain the desired representation (i.e., feature extractor) by pruning the output layer as illustrated in Fig. 1 A, resulting in a feature vector *Z* of size *N* = 32.

Our approach involves two data sets denoted *X*_*source*_ and *X*_*target*_. The source data set *X*_*source*_ consists of simulated sub-tomograms and corresponding class labels. These labels are automatically obtained from the simulation algorithm and therefore do not require manual annotation. The data set *X*_*source*_ is used to learn the representation *f*_*θ*_. On the other hand, we have the target data set *X*_*target*_ which consists of real sub-tomograms. This data set is not involved in training whatsoever. We also have labels for *X*_*target*_ which are only used to evaluate the clustering achieved on *f*_*θ*_(*X*_*target*_).

## 4 Experimental results

In this section, we first describe the data sets *X*_*source*_ and *X*_*target*_ and our training procedure, then we define our evaluation metrics, and finally present our results.

### 4.1 Description of synthetic and real data sets

#### Source data set

To our knowledge, SHREC is the only publicly available benchmark for evaluating algorithms on localization and classification tasks in cryo-ET. This data set is based on a physics-based model and is constituted of 10 tomograms with 10Å/voxel resolution and a size of 512 *×* 512 *×* 512 voxels. The tilt range is *±*60° with a tilt increment of 2°. In the benchmark published in 2021 [19], each tomogram consists of up to 1500 macromolecules, as well as membranes and gold fiducials. The macromolecule population is constituted of 13 different classes (obtained from the Protein Data Bank) of varying molecular weights (from 42kDa to 3326kDa). This data set has been designed to detect the break-point of particle picking algorithm, and contains several macromolecule classes which have very low weights (5 classes are below 200kDa).

From these tomograms, we extract a set of sub-tomograms of size 40 *×* 40 *×* 40 voxels, resulting in a total amount of 12905 training sub-tomograms and 1189 validation sub-tomograms.

#### Target data set

Our target data set is obtained from a set of real tomograms depicting *Chlamydomonas Reinhardtii* cells. The tomograms have a 13.68Å/voxel resolution and a size of 928*×*928*×*464 voxels. The tilt range is *±*60° with an increment of 2°. This data set was used in several studies [20, 21, 22, 23, 2] and was carefully annotated by experts. We gathered annotations for membrane-bound ribosomes, cytoplasmic ribosomes (≃ 3.2 MDa), proteasomes (≃ 750 kDa) and rubisco complexes (≃ 560 kDa). These annotations were exploited to extract the corresponding sub-tomograms of size 40 *×* 40 *×* 40 voxels. We resized the sub-tomograms to obtain a 10Å/voxel resolution to fit the source data set. In the end, our test set is constituted of 500 sub-tomograms (100 per class).

For both data sets, we also collected sub-tomograms to constitute the negative class. In order to make sure that these do not contain macromolecules, they were sampled outside of the lamella. An illustration of both data sets is shown in Fig. 4.

### 4.2 Training procedure

For training we used the ADAM algorithm, chosen for its good convergence rate, by setting the learning rate to 10^*−*4^, the exponential decay rate to 0.9 for the first moment estimate and to 0.999 for the second moment estimate. We used the categorical cross-entropy as training objective and set the batch size to 64.

The number of examples per class varies slightly in the SHREC data set. Therefore, we apply the same resampling strategy as in [2], so that the distribution of classes in a batch is uniform. In order to speed up the training, we initialized the network with weights obtained from the DeepFinder network, trained for the SHREC 2021 challenge [19].

### 4.3 Methods and metrics for visual assessment and classification evaluation

We evaluated our unsupervised proof-of-concept method on the validation and test data sets described in Section 4.1.

For a visual assesment of the learned data representation, we embedded the feature space in ℝ^2^ using the t-SNE algorithm [24]. T-SNE is a non-linear dimensionality reduction technique, whose objective is to find a faithful low-dimensional representation of a high-dimensional space. We obtained similar results by appying UMAP [25] for visual assessment.

Next, we evaluated the k-means clustering in terms of homogeneity, completeness and V-measure. Those metrics are based on normalized conditional entropy measures of the clustering with respect to ground truth labels. The values range from 0 to 1. Values close to 1 correspond to a better classification. Homogeneity measures the ratio of samples of a single class pertaining to a single cluster. Completeness measures the ratio of members of a given class that are assigned to the same cluster. The V-measure is the harmonic mean between homogeneity and completeness.

### 4.4 Evaluation of the results

First, we assess the learned representation on *X*_*source*_ (see Fig. 2 A,C). In the two-dimensional space calculated by the t-SNE algorithm, the classes are organized in well separated groups, with very few data points belonging to the wrong groups. In order to test if the representation *Z* is clustering friendly, we apply k-means and compute metrics for different parameter values (i.e. the number of clusters). The best V-measure (0.96) is obtained when the cluster number parameter is 13, which corresponds to the number of classes.

**Figure 2:**
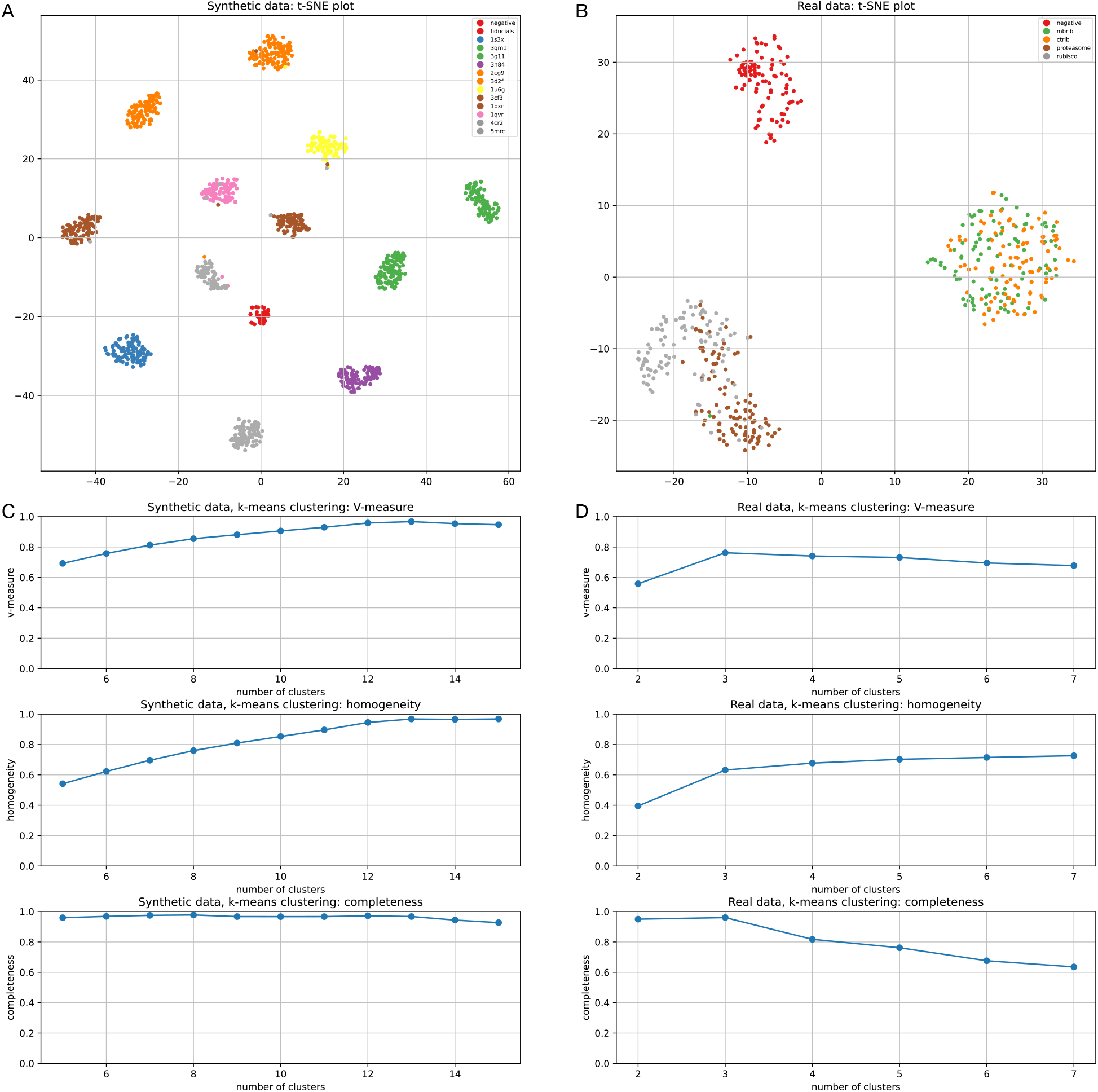
Evaluation of the learned representation. (A,B) 2D embedding of the 32-dimensional feature space. The embedding is obtained with the t-SNE algorithm [24]. We illustrate embeddings for (A) the source data (simulated, SHREC’21) and for (B) the target data (real, *Chlamydomonas Reinhardtii* cells). (C,D) For each data set we applied k-means clustering in conjunction with the representation, and computed the V-measure, homogeneity and completeness with respect to the number of clusters (the parameter of k-means). All results were obtained on the test data.

Next, we tested the generalization potential of *f*_*θ*_ on *X*_*target*_. As is illustrated in Fig. 2 B, the t-SNE plot shows that the data points are organized in three main groups: one group corresponds to the negative class, another one to the membrane-bound and cytoplasmic ribosomes, and a last one to the proteasomes and rubisco complexes. Accordingly, this feature space allows to discriminate well ribosomes and the negative class. Note that this representation has difficulties differenciating *mb-ribos* from *ct-ribos*. This can be explained by the fact that SHREC does not contain examples of membranebound macromolecules (altough it contains membranes). Therefore the network learned to focus the classification on the 3D structure (as expected), but without taking the neighborhood into account. Yet the context is precisely what differenciates membrane-bound from cytoplasmic ribosomes. This plot also shows that the representation does not separate well the proteasome and rubisco complex classes. That being said, we observe two sub-groups, each with an over-representation of one of these classes.

According to Fig. 2 D, the best V-measure (0.76) is obtained when the cluster number is set to 3. In this case, homogeneity shows that each cluster contains only members of a single class in 63% of the cases, and completeness shows that in 96% of the cases all members of a given class are assigned to the same cluster. We note that the same completeness score as for the source data set has been obtained, but the homogeneity is significantly lower (0.63 instead of 0.96). The homogeneity measure is consistent with the visual analysis provided by t-SNE, and this value drop demonstrates the limited ability to transfer *f*_*θ*_ from synthetic to real data.

Nonetheless, while these results point out the weaknesses of this approach, it is also clear that *f*_*θ*_, which has been obtained without manual annotations, is able to structure real data. If we examine the composition of the clusters in Fig. 3 B when the cluster number is set to 4, cluster #1 contains 100% of the negative sample class, cluster #2 contains 98.5% of ribosomes (regardless of their binding state), cluster #3 contains 73% of proteasomes and cluster #4 contains 84% of rubisco complexes. These results are still preliminary but are promising, leading us to believe that deep learning based unsupervised classification methods will play an important role in the analysis of cryo-ET data in the forthcoming years.

**Figure 3:**
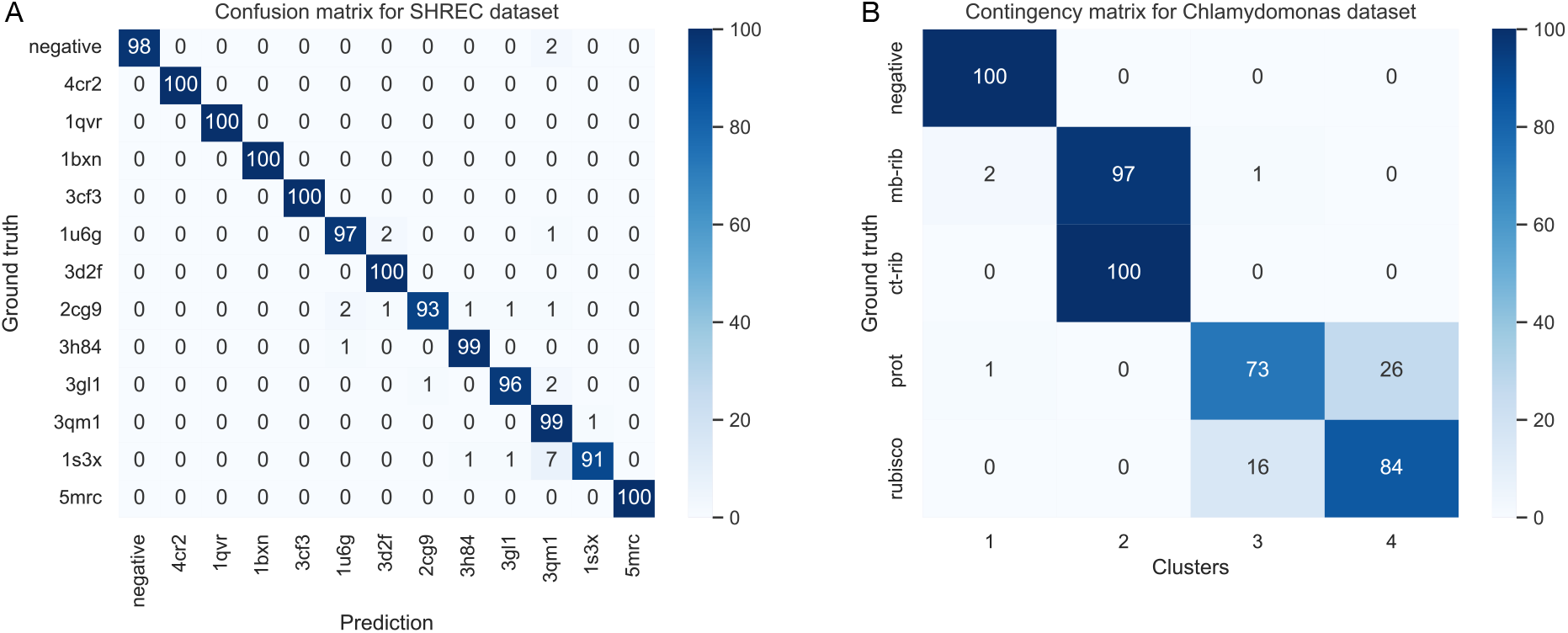
(A) Confusion matrix of the classification (supervised) on the SHREC dataset. (B) Contigency matrix of the k-means clustering (unsupervised) on the Chlamydomonas dataset. Displayed values are normalized and correspond to percentages.

**Figure 4:**
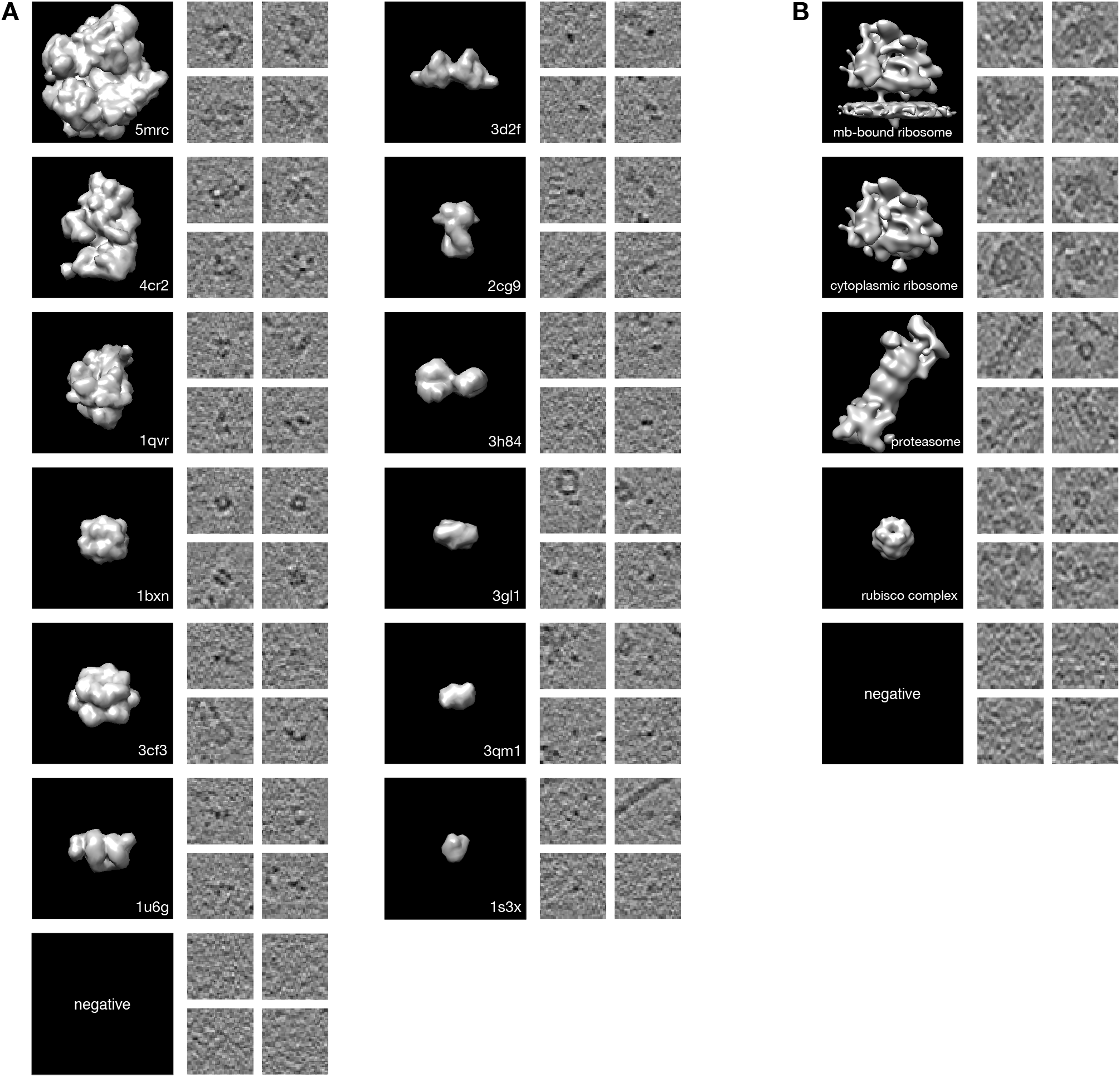
Dataset illustrations. (A) Synthetic dataset SHREC’21 and (B) Real dataset (Chlamydomonas Reinhardtii). For both datasets, we display the macromolecule classes. For each class, on the left side a 3D view of the pdb structure, and on the right four instances of this class as found in the dataset. The instance images correspond to the central *xy* slice of the 3D volume. For enhancing the contrast, the slice has been averaged with the two neighboring slices in each direction.

## 5 Discussion and conclusion

In this paper, we provided a brief overview of unsupervised classification methods based on deep learning. We presented the challenges, obstacles and promises to analyze cryo-ET data.

Our experiments suggest that an adapted strategy is to decouple representation learning from the clustering (as opposed to deep clustering methods). Learning the representation from a simulated source data set allows one to better control what the network encodes. This facilitates interpretability of the feature space, which is an important asset in the task of studying unknown macromolecules. As such, we have succeeded in obtaining a desired feature of this representation, namely an invariance to the macromolecule orientation, which has been achieved by design of the source data. Rotation invariance is an appealing property for reducing computation time, as it avoids the use of sub-tomogram alignment procedures, as is the case with conventional methods [3, 4, 5]. Moreover, this approach is very fast as it only took 8 seconds to classify 400 sub-tomograms of size 40 *×* 40 *×* 40 on a Tesla K80 GPU. On the other hand, training took 2 hours, which is relatively short for deep learning. That being said, the advantage of the proposed approach is that the user does not need to train the network himself.

This approach can be improved in future works. First, we can improve the definition of the source data set. SHREC’21 is composed of 9 training tomograms (12905 sub-tomograms) and 13 classes, which is small compared to benchmarks like ImageNet. As a good data model is available, we can potentially generate infinite amount of data to increase the size of the training set. Also, we can increase the number of macromolecule classes. In particular, we can add different conformation states of the same macromolecule species, as well as several binding states (e.g., membrane-bound). Training on such a data set would force the network to encode a more detailed representation of the macromolecule structure and its biological local context. We showed that currently, differentiating binding-states is one limitation of this approach.

Second, we would like to point out that our results have been achieved with under-sampled data, i.e. a pixel size of 10Å, while methods like [3, 4, 5] use full resolution. In fact, the results presented in [3] and [4] have been obtained with a pixel size of 6Å, and in [5] with a pixel size of 2.62Å. Consequently, reducing the voxel size of the source data set represents one axis of improvement for this strategy.

Finally, there is a generalization issue when switching from the source to the target data set. This low performance results from the fact that the source and target data do not have the same statistical distribution. The feature extractor *f*_*θ*_ has been fitted to the source distribution only, and hence the clustering performance on the target distribution is sub-optimal. The family of methods that aim to address this issue is known as *domain adaptation*. Here, the focus is on learning a representation that is both discriminative and invariant to the change in data distributions [26, 27]. Recently, such methods have been applied successfully to cellular electron microscopy images [28, 29]. We believe that domain adaptation constitutes a promising perspective to achieving robust unsupervised classification for cryo-ET.

## Acknowledgements

This work was jointly supported by the Fourmentin-Guilbert Foundation and Region Bretagne (Brittany Council). Computing was performed on the Inria Rennes computing grid facilities partly funded by France-BioImaging infrastructure (French National Research AgencyANR-10-INBS-04-07, Investments for the future).

We thank B. D. Engel, T. Peng, L. Lamm, R. D. Righetto, W. Wietrzynski, S. Pfeffer and S. Albert for providing data sets with annotations (ribosomes [20], proteasomes [21] and rubisco complexes [22]), expertise and fruitful discussion.

We also thank E. Fourmentin, D. Lariviere, J. Ortiz and W. Baumesiter for helping design research. We thank A. Martinez for his critical feedback and valuable suggestions about machine learning with synthetic datasets in cryo-ET.

Finally, we thank the organizers of the SHREC challenges [19] for helpful assistance: I. Gubins, R.C. Veltkamp, G. van der Schot and F. Forster.

## Data availability

The synthetic dataset is available on the website of the SHREC 2021 challenge (http://www2.projects.science.uu.nl/shrec/cet/). Tomograms from the real dataset of *C. reinhardtii* cells can be found in the EMDB under accession numbers EMD-3967 and EMD-12749.

## Notes

### Competing Interest Statement

The authors have declared no competing interest.

